# Estimating the Mutational Fitness Effects Distribution during early HIV infection

**DOI:** 10.1101/185678

**Authors:** Eva Bons, Frederic Bertels, Roland R Regoes

## Abstract

The evolution of HIV during acute infection is often considered a neutral process. Recent analysis of sequencing data from this stage of infection, however, showed high levels of shared mutations between independent viral populations. This suggests that selection might play a role in the early stages of HIV infection. We adapted an existing model for random evolution during acute HIV-infection to include selection. Simulations of this model were used to fit a global mutational fitness effects distribution (MFED) to sequencing data of the *env* gene of individuals with acute HIV infection. Measures of sharing between viral populations were used as summary statistics to compare the data to the simulations. We confirm that evolution during acute infection is significantly different from neutral. The distribution of mutational fitness effects is best fit by distribution with a low, but significant fraction of beneficial mutations and a high fraction of deleterious mutations. While most mutations are neutral or deleterious in this model, about 5% of mutations is beneficial. These beneficial mutations will, on average, result in a small but significant increase in fitness. When assuming no epistasis, this indicates that at the moment of transmission HIV is near, but not on the fitness peak for early infection.

## Introduction

Evolution is driven by new mutations causing a change in fitness of the organism. If a new mutation increases the reproductional success, or the fitness, this mutation is likely to be selected for and eventually fix in a population. However, most mutations are not beneficial to the organism. Instead they are neutral - having no effect on the fitness, or deleterious, reducing the amount of offspring compared to the ancestor. The effects a mutation can have on the fitness lie on a continuum from completely lethal to beneficial, including viable but deleterious effects and neutral effects. The mutational fitness effects distribution (MFED, reviewed by Eyre-Walker and Keightley, 2007) captures how these effects are distributed for a certain organism in a certain environment.

The MFED has been inferred for several viruses using site-directed mutagenesis studies (Sanjuán, 2010). It has a similar shape across the different virus species, with a sizable amount of mutations (20 to 40%) being lethal and the rest forming a single peak at or close to zero. In some, but not all viruses, there is a small amount (less than 10%) of beneficial mutations. Knowing the MFED of an organism in a certain environment can help us understand and predict the evolutionary dynamics of this organism in this environment.

Early HIV infection is characterized by rapid expansion of the virus population. Transmission is a bottleneck, as only one to five viruses are responsible for the establishment of infection (Keele et al., 2008). After a few weeks, virus levels can reach up to 10^6^ virus particles per ml plasma (Fiebig et al., 2003). This expansion is accompanied by rapid sequence diversification due to the high mutation rate of HIV. This rate is estimated to be between 1.1 · 10^−5^ to 5.8 · 10^−3^ mutations per base per replication, depending on the method and source material of the estimation (Mansky and Temin, 1995; Huang and Wooley, 2005; Dapp et al., 2013; Cuevas et al., 2015).

During these early stages of infection, HIV evolution is often considered a neutral process due to the rapid expansion and absence of immune response, which is the main evolutionary pressure on HIV during infection. Keele et al. (2008) and Lee et al. (2010) showed that sequence patterns in the *env*-gene of 82 individuals show typical signs of neutral evolution during acute infection, such as a star-like phylogeny. While the mutational patterns appear neutral on an individual level, signs of selection become clear when looking at the viral populations in several individuals at once. Wood et al. (2009) and Bertels et al. (2017) studied the same dataset and found several convergent mutations; mutations that appear in several viral populations independently. These mutations are likely positively selected, indicating evolution during early HIV infection is not neutral.

By estimating the MFED for HIV during early infection, evolutionary pressures during these early stages of infection can be better understood. There have been several attempts at estimating fitness effects of mutations for HIV, but they either only consider the amino-acid level (Haddox et al., 2016; Ferguson et al., 2013) or analyze a subset of mutations only, such as deleterious mutations (Zanini et al., 2017) or resistance mutations (Martinez-Picado and Martínez, 2008).

In this paper, we use these patterns of convergent evolution found by Wood et al. (2009) and Bertels et al. (2017) to estimate the MFED of HIV during early infection. For this, we use simulations of sequence evolution including selection that were fitted to patterns of sharing in *env*-sequences collected by Keele et al. (2008) and Li et al. (2010).

## Results

### Simulation of the molecular evolution of HIV during early infection

To investigate the impact of viral fitness differences on HIV sequence evolution in infected individuals we developed a simulation model of early HIV infection. The model is based on the Monte Carlo simulations of the synchronous infection model for HIV sequence evolution presented in Lee et al. (2010). In contrast to Lee et al., however, we relax the assumption that mutations are neutral. Instead, we assign a fitness advantage or disadvantage to each mutation that occurs according to a mutational effects distribution. We parameterized this distribution fairly flexibly, such that it describes a fraction of beneficial, detrimental and completely lethal effects, and can also be collapsed, for appropriate parameters, to a fully neutral model (as Lee et al.).

A simulation starts with a randomly generated sequence with a relative fitness of one. In every generation, all sequences in the population generate N new offspring, where N is a random draw from a Poisson distribution with mean *R*_0_·*ƒ*. In this formula, *R*_0_ denotes the absolute fitness of the virus, which we set to 6 in accordance with Lee et al. (2010), and *ƒ* the relative fitness compared to the ancestor sequence. Point mutations can occur in these offspring sequences, which can alter the fitness of the sequence.

Our goal with these simulations is to generate datasets of the same structure as the dataset by Keele et al. and Li et al. by matching the number of generations in our simulation to the time since infection of the infected individual in the empirical datasets. All known estimates for the time since infection, however, are based on a neutral model of evolution, which can lead to a bias. To avoid this potential bias in the time since infection, simulations were run for as long as necessary to match the amount of unique mutations in a sample of the same size as available in the data set. The resulting sequence sample (see figure 1) has similar mutational characteristics as the sequence sample available for each subject, without requiring to specify for how many generations the simulation should be run.

**Figure 1:**
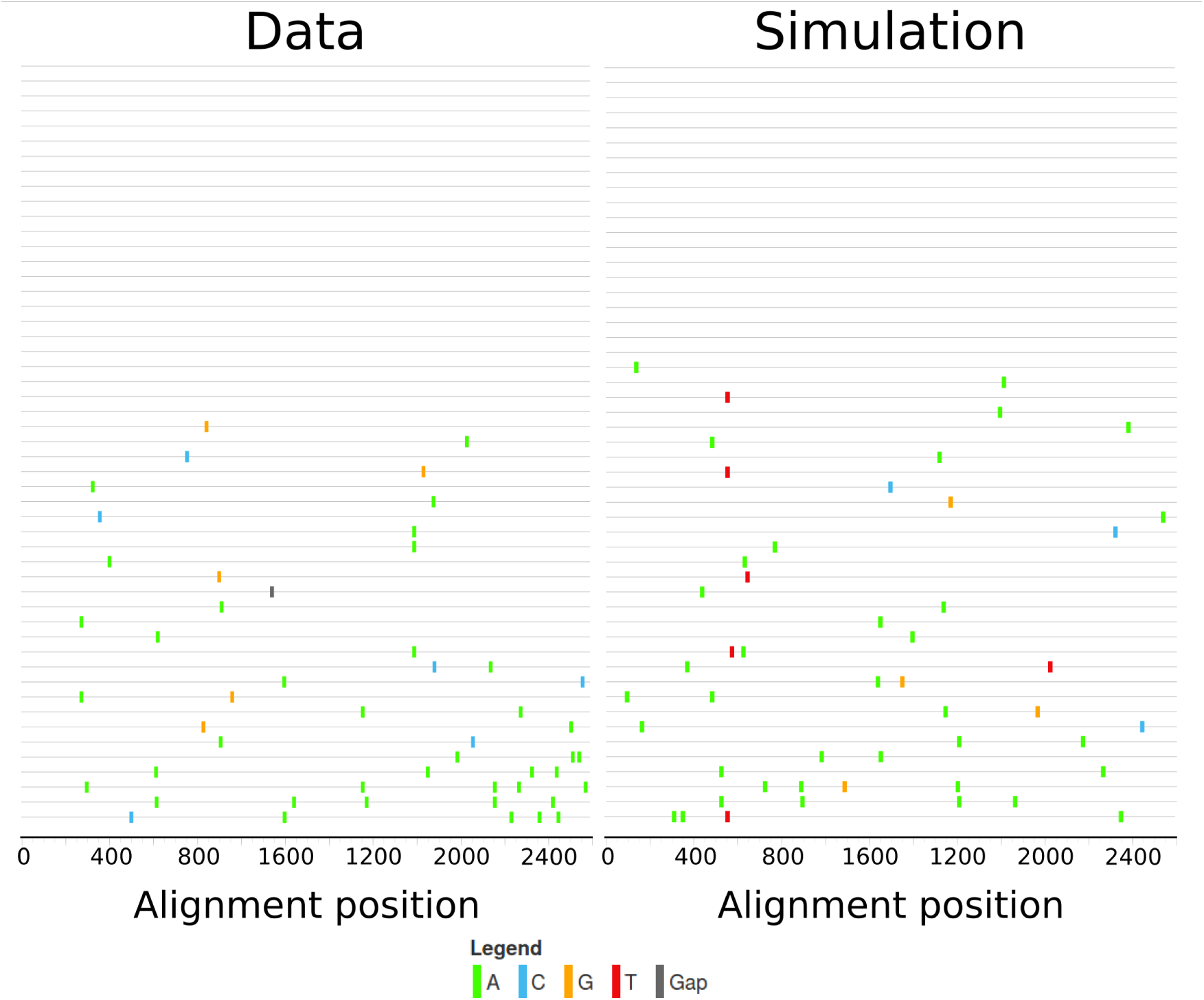
visualization of sequence samples using highlighter (hiv.lanl.gov) of the sequence sample from subject 1018 (left) and the output of a simulation matching this data (right). Both sequence samples consist of 51 sequences. In the empirical sample, there are 50 mutations, of which 44 occur only in a single viral population. 24 sequences are unmutated. In the simulated sample, there are 49 mutations, of which 44 occur only in a single viral population and 20 sequences are unmutated

### Estimating the shape of the mutational effects distribution

In order to calculate the fitness of a mutated sequence, every possible mutation is assigned a fitness effect according to the mutational fitness effects distribution (MFED). Each of these effects is assumed to apply universally in all hosts, there are no host-specific effects. The fitness of a sequence is then the product of the fitness effects of all mutations in the sequence.

The effects in the MFED range from zero to infinity, with a fitness effect of one indicating a neutral mutation (see figure 2). Deleterious mutations have a fitness effect smaller than one, with an effect of zero indicating a lethal mutation. Sequences carrying a lethal mutation differ from sequences carrying a non-lethal deleterious mutation since they will never produce any offspring, while sequences carrying mutations with a very small, but not 0, fitness effect can still produce offspring, albeit with a very low probability. A very beneficial mutation might also compensate for such a deleterious mutation, while this is impossible in the case of lethal mutations.

**Figure 2:**
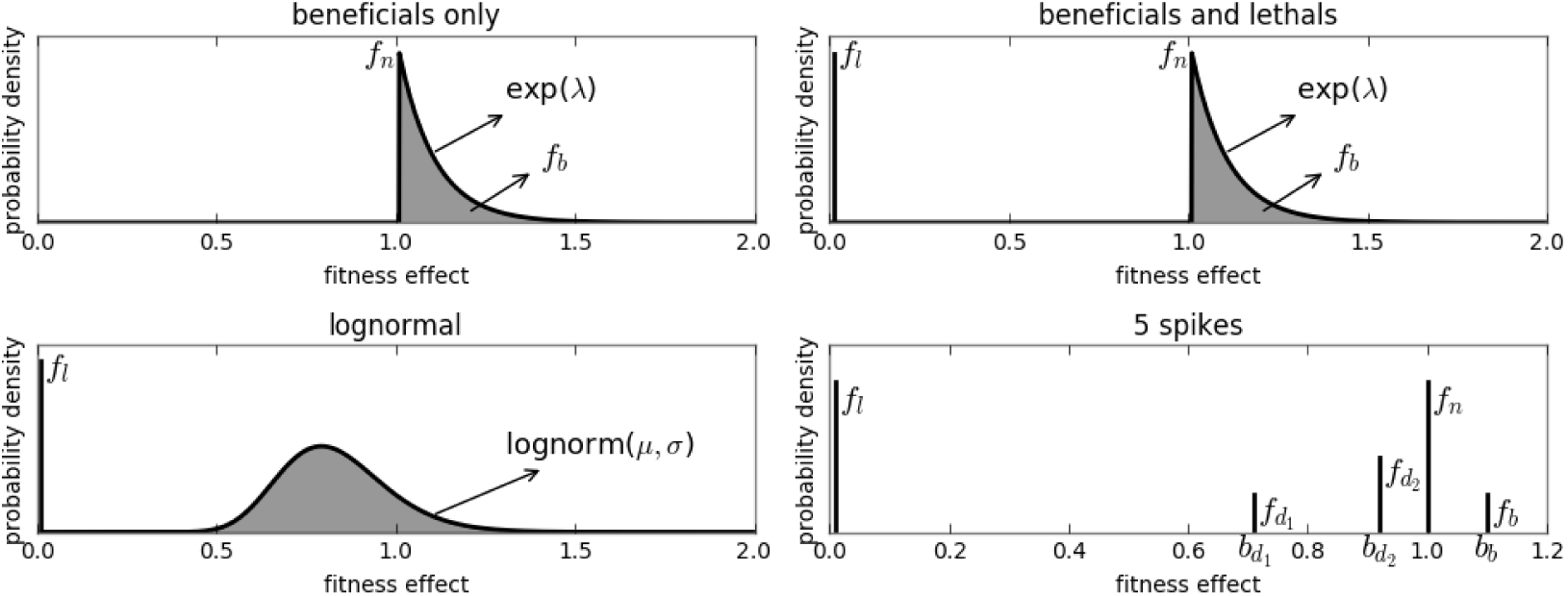
Overview of 4 of the 6 different models for the mutational fitness effects distribution (MFED). The beneficials only model has 2 free parameters (*ƒ*_*b*_,λ), the beneficials and lethals model has 3 free parameters (*ƒ*_*l*_,*ƒ*_*b*_, λ), the log-normal model has 3 free parameters (*ƒ*_*l*_,*μ*, *σ*). The 5 spikes model has 7 free parameters (*ƒ*_*l*_, *ƒ*_*d*_1__, *b*_*d*_1__, *ƒ*_*d*_2__, *b*_*d*_2__, *ƒ*_*b*_, *b*_*b*_)

The fitness effects distribution will affect the amount of shared mutations across viral populations. While the probability of a mutation occurring does not change, the probability of a mutation being maintained in the population and later sampled is affected by the fitness effect of the mutation. Once a beneficial mutation occurs, the sequence carrying this mutation will create more offspring than unmutated sequences, and will be overrepresented in the viral population after a few generations. This increases the chance of observing the mutation in two or more samples.

Lethal or deleterious mutations will cause the sequence carrying the mutation to have fewer offspring and are therefore unlikely to be observed. Since these mutations are so unlikely to be observed, the sites can be considered immutable, which results in an effective shortening of the genome available for mutation. This might indirectly increase the chance of sharing mutations by increasing the chance that other, less detrimental, mutations are observed.

We defined 6 different models for the mutational effects distribution (see figure 2), additional to a neutral model where all fitness effects are one.

The first two models describe simplified distributions with a restricted effects range. The ‘beneficials only’ model consists only of neutral and beneficial fitness effects, and is defined by a fraction of beneficial mutational effects (*ƒ*_*b*_), which are exponentially distributed with mean λ. The rest of the mutational effects are neutral (*ƒ*_*n*_ = 1 – *ƒ*_*b*_). The second model, ‘lethals only’, consists of a fraction of lethal (*ƒ*_*l*_, fitness effect of 0) and neutral effects (*ƒ*_*n*_ = 1 – *ƒ*_*l*_). The third model is a combination of these, whith a fraction of lethal mutations, a fraction of exponentially distributed beneficial mutations and the rest of the mutations is neutral. The last three models are more complex and try to capture a wider spectrum of conceivable fitness effects and replicate the observed distributions in other organisms mentioned in the introduction. The log-normal model consists of log-normally distributed fitness effects, with parameters *μ* and *σ*, in the entire range from zero to infinity, plus a certain amount of fitness effects that are exactly 0, the fraction lethals (*ƒ*_*l*_). A variation of this model is the ‘log-normal truncated’ model, which is the same as the log-normal model, only there are no beneficial mutations. Instead, all mutations that would have been beneficial in the log-normal model are now neutral. The last model, ‘5 spikes’, allows for exactly five different values for the fitness effects. Fitness effects can be 0 (lethal), 1 (neutral), one of two values (*b*_*d*_1__ and *b*_*d*_2__) between zero and one (deleterious) or a value larger than one (*b*_*b*_, beneficial). The relative amount of mutations with each effect are represented as *ƒ*_*l*_,*ƒ*_*d*_1__,*ƒ*_*d*_2__,*ƒ*_*n*_ and *ƒ*_*b*_. This allows for an approximation of multi-modal models without more complex model definitions. The neutral model, where all mutations have a fitness effect of one is recovered by setting all parameters to zero in any of the models.

We use ABC-SMC (Approximate Bayesian Computation using Sequential Markov Sampling) to do simultaneous model selection and parameter estimation. The SMC procedure start with a set of parameters for all models, where all models are equally represented. For each parameter set a simulation is performed and those simulations whose distance to the data is below a threshold are accepted. In each subsequent iteration the threshold is lowered and new parameter sets are sampled from the set of accepted parameters in the previous iterations. This results in simulataneous parameter estimation and model selection, since a good model will have more parameter sets accepted, where bad models will have less accepted parameter sets. If no parameter sets are accepted for a certain model, this model has ‘died out’ and is considered a very unlikely model candidate.

This procedure requires summary statistics to calculate the distance between simulations and the data. The summary statistics used here (see *methods* for a full overview) are based on measures of sharing, such as the distribution of the average degree of sharing of the mutations in a viral population and the number of populations the mutations appear in, but also population-specific statistics such as inter-sequence hamming distance and the number of unmutated sequences in the sample were used.

### The best estimate for the MFED

The model selection results in two equally likely models: the ‘lethals and beneficials’ model and the log-normal model. A summary of the parameter estimation and characteristics of the MFED with best fit parameters for these models can be found in table 1. Both models have a relative probability (‘support’) of approximately 0.4. The ‘beneficials only’ and ‘5 spikes’ model have a much lower support, both are less than 0.1. The neutral model, ‘lethals only’ model and the truncated log-normal model died out before the 4th iteration of the ABC-SMC.

**Table 1:** fitted parameters with 95% HPD and moments of the MFED for the top-2 models

|  | **lethals and beneficials** | **log-normal** |
| --- | --- | --- |
| parameters | $f_l=0.182(0.019;0.552)$<br>$f_b=0.053(0.008;0.228)$<br>$\beta=0.202(0.051;0.732)$<br>$G \rightarrow A=10.451(2.361;69.169)$ | $f_l=0.045(0.003;0.422)$<br>$\mu=-0.248(-0.284;-0.088)$<br>$\sigma=0.149(0.104;0.288)$<br>$G \rightarrow A=10.621(1.886;74.797)$ |
| support | 0.439 | 0.409 |
| mean effect | 0.828 | 0.753 |
| fraction beneficial | 0.052 | 0.045 |
| beneficial effect | 1.204 | 1.065 |
| beneficial gain | 0.063 | 0.048 |
| fraction neutral | 0.766 | 0 |
| fraction deleterious | 0 | 0.909 |
| deleterious effect | 0 | 0.776 |
| fraction lethal | 0.182 | 0.046 |
| deleterious loss | 0.182 | 0.250 |

For the top-2 models, enough simulations were accepted to use for parameter estimation. Many parameters have a relatively large 95% highest probability density (HPD) interval, although all parameters are significantly different from the neutral value. The MFEDs corresponding to the best fit models are very similar. Both distributions have a mean fitness effect of approximately 0.8. The log-normal model has slightly fewer beneficials (4.5% vs 5.2% in the ‘lethals and beneficials’ model), that also have a slightly lower effect. It is harder to compare the deleterious and lethal mutations between the two models, since the ‘lethals and beneficials’ model does not include deleterious mutations, while in the log-normal model, 90% of mutations is deleterious, albeit with a low effect. The deleterious loss (the product of the fraction of deleterious/lethal mutations and their loss of fitness, which equals 1 minus the fitness effect) is the easiest way to compare them. Although the ‘lethals and beneficials’ model has more lethal mutations, 90% of mutations in the log-normal model are deleterious, leading to an overall higher loss due to deleterious and lethal mutations.

In figure 3, two measures of sharing are compared between the data and the simulations using different models for the MFED: the number of viral populations each mutation appears in, and the degree of sharing; the average amount of populations the mutations in one population are shared with. The simulations from the best-fit models resemble the data much better than the neutral simulations, although the distributions are not perfectly recovered. Interestingly, the log-normal model - which has more deleterious and less beneficial effects results in more convergent evolution than the ‘lethals and beneficials’ model. In the former, the distributions for both measures are shifted to the right compared to the data, indicating higher amounts of sharing, while in the latter, the distribution is slightly shifted to the left compared to the data.

**Figure 3:**
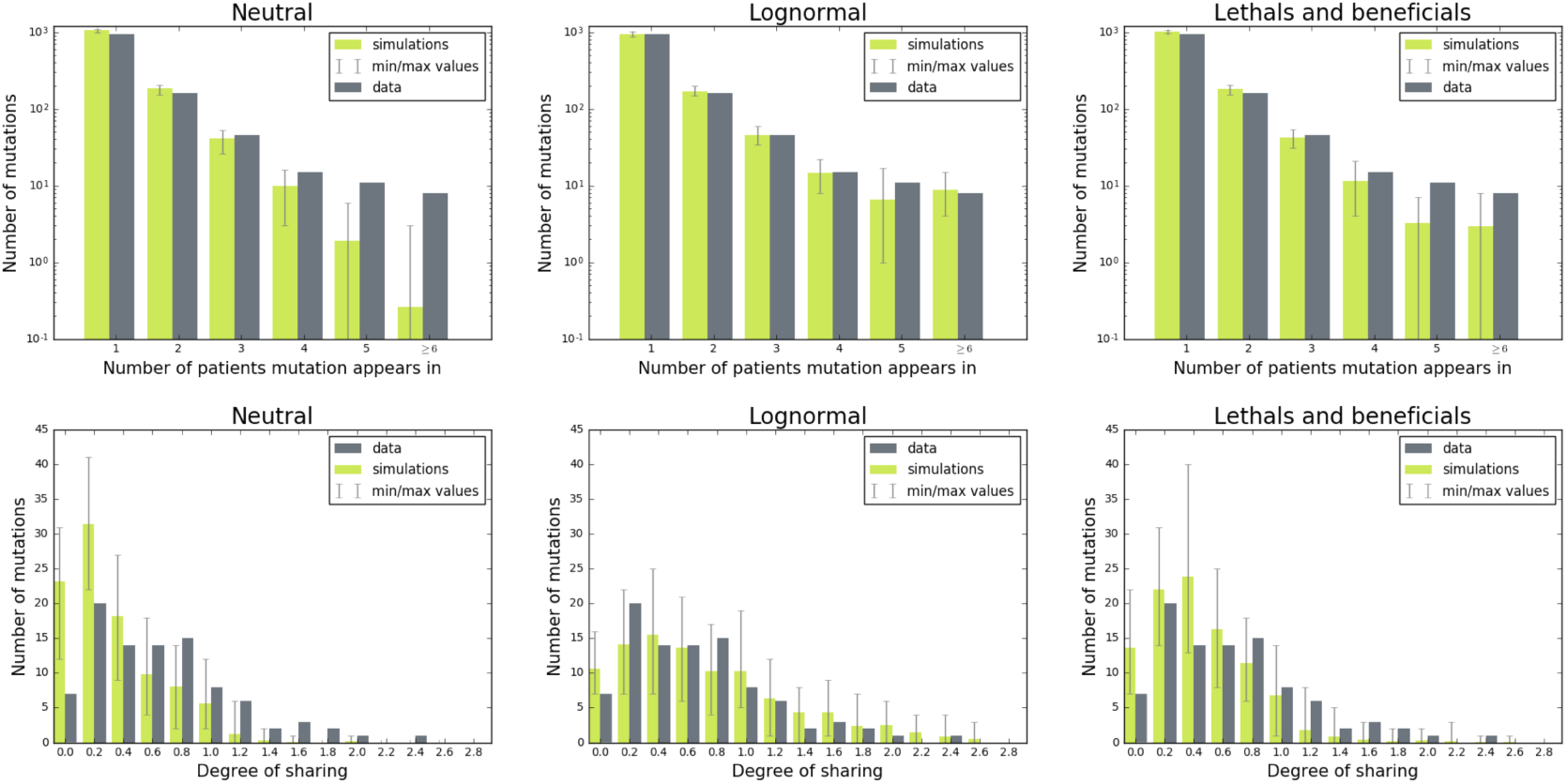
Measures of sharing in the data compared to the mean of 100 simulations: the number of populations each mutation appears in (top) and the degree of sharing (the average amount of populations the mutations in one subject are shared with, bottom) left: simulations where mutational effects are neutral, center: simulations where mutational effects are distributed according to the log-normal model, right: simulations where mutations are distributed according to the ‘lethals and beneficials model

### Fitness of shared mutations

Having an estimate of the MFED and being able to reproduce the datasets with this MFED allows us to estimate the fitness effect of the shared mutations. This can be achieved by finding the average fitness effect a mutation in a certain frequency class has (see figure 4). For both models, mutations found in more than four populations have a MFED with a median higher than one, indicating that this mutation is likely beneficial. The average effect of these mutations differs between the models. If the MFED is distributed according to the ‘lethals and beneficials’ model, the fitness effect of a mutation occurring in 14 populations has a fitness effect between 2 and 3 (indicating this mutation will cause 2 to 3 times as much offspring per generation). The effect of such a mutation is much lower when the MFED is distributed according to the log-normal model: the effects there are then between 1.2 and 1.4.

**Figure 4:**
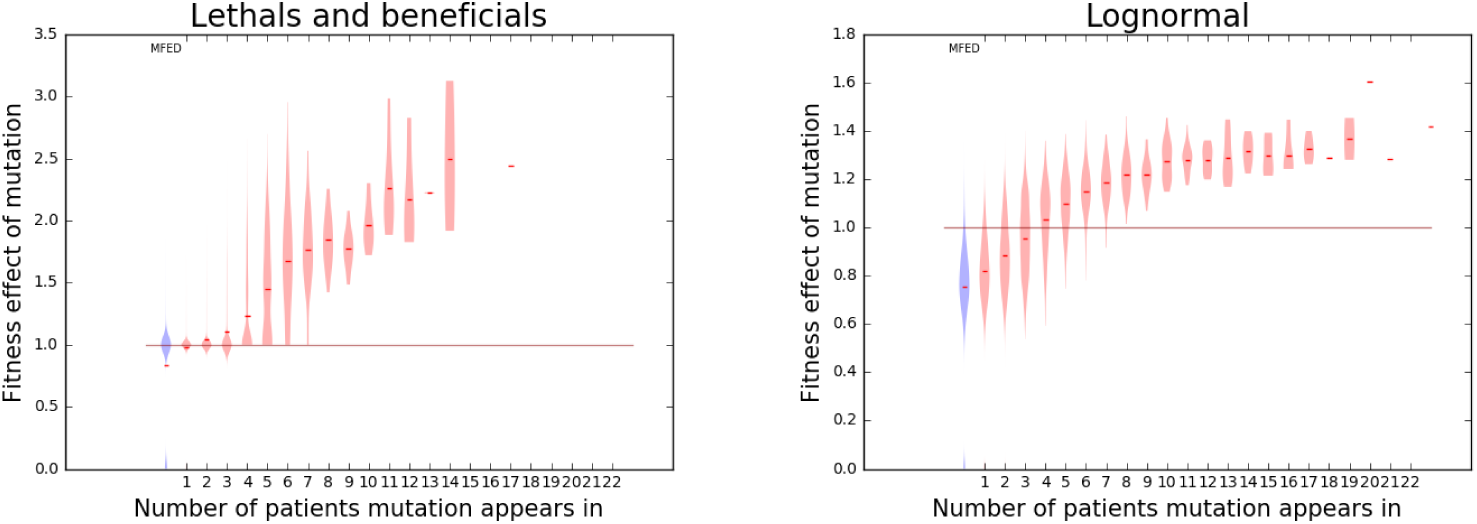
Violin plot representing the distribution of fitness effects per frequency class, derived from 100 simulations using best-fit parameters for both models. The blue violin on the left represents the original MFED, the red lines indicate the median fitness effect in the respective frequency class

### The role of APOBEC

The mutagenic enzyme APOBEC left a detectable imprint in the data by Keele et al. and Li et al. Mutations carrying an ‘APOBEC signature’ make up 39% of the shared mutations in the dataset, while they only make up 12% of all occurring mutations. 19% of populations carry APOBEC-mediated mutations. In these populations, 45% of G-to-A mutations are APOBEC mediated. The distribution of the degree of sharing per subject for the APOBEC populations is significantly higher from the populations without any APOBEC mutations (t-test, p-value 4.74 · 10^−4^).

Since the effect of APOBEC-mdiated hypermutation has an effect on the patterns of sharing, it is important that we include this increase in mutation rate in the model fits and parameter estimates. For the simulations of all populations that carried APOBEC-mediated mutations, the G-to-A mutation rate was increased.

Since estimates of the contribution of APOBEC to the mutation rate vary widely, we estimated the increase in the G-to-A mutation rate (*μ*_*GA*_*APOBEC*__ /*μ*_*GA*_) along with the parameters of the MFED. Initially, these estimates were all on the lower end of the prior distribution, which ranged from 1-160, while the 95% HPD of the parameter estimate across models ranged from 1 to 34. To make sure the G-to-A mutation rate increase is significantly different from one, we re-estimated the parameters of the log-normal model with a logarithmic prior distribution for *μ*_*GA*_*APOBEC*__ /*μ*_*GA*_ from *e*^−10^ to *e*^5^. The estimates remained the same, but the 95% HPD intervals no longer included one, indicating a significant increase in the G-to-A mutation rate due to APOBEC.

## Discussion

The combined dataset of Keele et al. and Li et al., containing sequencing data of the *env*-gene from over a hundred inidividuals in early stages of HIV infection, can provide many insights into the dynamics of HIV at or closely after the moment of infection. The conclusions made by Keele et al. (2008), Li et al. (2010), and Lee et al. (2010) about the number of founder viruses and the time since infection, are all based on an assumption of neutral evolution. However, Wood et al. (2009) and Bertels et al. (2017) found evidence for positive selection during early HIV infection using this dataset.

We fitted a global mutational effects distribution to sequencing data from Keele et al. (2008) and Li et al. (2010), where mutations are assumed to have the same effect in all inidividuals, independent of time, environment or sequence context. We neglect epistasis in our model, since we only consider early infection where the number of mutations per sequence is expected to be low. Interactions between mutations are therefore unlikely.

The fit resulted in two high-probability models (the ‘lethals and beneficials’ and the log-normal model), two low-probability models (the ‘beneficials only’ and ‘5 spikes’ model), and three exceedingly unlikely models (neutral, ‘lethals only’ and the truncated log-normal model). Although we cannot distinguish between the top two models, their shared characteristics allow us to make several conclusions about the MFED. Both models contain a small fraction of beneficial mutations, indicating that the high amount of sharing cannot be only due to the effect of deleterious mutations, which reduces the chance of observing these mutations, thereby increasing the chance to observe other, less deleterious mutations. However, the presence of more beneficial mutations does not necessarily mean more sharing, as becomes apparent in the log-normal model. Here, the high amount of deleterious mutations causes higher levels of sharing than the ‘beneficials only’ model, even though this model contains less beneficial mutations. The presence of deleterious mutations seems to reduce the amount of beneficial mutations needed to reach the same level of shared mutations between populations. This observation, together with the poor performance of models which do not include deleterious mutations highlights the importance of deleterious mutations when studying convergent evolution on the sequence level.

Due to its high flexibility, we expected the ‘5 spikes’ model to reach higher model probabilites. We suspect that this is not the case because of the high dimensionality of this model (8 parameters, vs 1-3 parameters for the other models), which is inherently punished by the ABC-SMC method (Toni et al., 2009). With higher sampling rates, it is possible that a good parameter set for this model can be found, that might perform even better than the current best models. However, the computational costs for this will be very high.

The log-normal estimate for the MFED is most in line with estimates in other viruses from single-nucleotide substitution studies (Sanjuán et al., 2004; Carrasco et al., 2007; Domingo-Calap et al., 2009). In these studies, beneficial mutations ranged from not present to 4% of occurring mutations, but small beneficial effects are difficult to detect in these studies (Eyre-Walker and Keightley, 2007). 20 to 40% of mutations were lethal and 30 to 50% deleterious. A quick calculation shows us that 4% of mutations will introduce a premature stop codon, which is typically lethal. This sets a lower bound on the number of lethal mutations, which is just met by the best fit of the log-normal model. If this model is correct, all lethal mutations in the *env*-gene would be due to premature stop codons. It is also important to note that these estimates are for a full genome, including non-coding regions. Our estimates only include the *env*-gene, which is under stronger evolutionary pressure than the rest of the genome (Zanini et al., 2015). We therefore expect that the MFED for this gene is more extreme than the full genome, leading to more beneficial and deleterious mutations.

This study also highlights the importance of APOBEC-mediated hypermutation in HIV evolution. The protein is part of the hosts’ innate immune response against viruses that mutagenizes single-stranded DNA (Goila-Gaur and Strebel, 2008) Considering that, according to the MFEDs estimated here, 20 to 95% of mutations will be deleterious or lethal, an increased amount of mutations will likely reduce the fitness of the virus, thereby limiting the viral spread. However, there is also a small chance of introducing beneficial mutations, which via selection will rapidly fix in the population. Selection of APOBEC-mediated mutations has been observed in HIV before (Kim et al., 2014; Wood et al., 2009) and our study suggests the same. Virus samples carrying APOBEC-mediated mutations show higher levels of sharing than those without APOBEC signature, which suggests that they contain more beneficial mutations due to the increased sampling of mutations caused by the higher mutation rate.

The concept of a MFED fails to account for a potentially changing fitness landscape due to changes in the environment. For example, fitness effects of mutations can depend on the composition of the viral population, i.e. a mutation may confer a benefit against some competitor strains but not others. This aspect is particularly pronounced in systems with non-transitive interactions (Chao and Levin, 1981). However, such interactions rely on the production of anti-competitor toxins that are less common in viruses.

Differences between hosts are also not described by a global MFED. While genetic differences between hosts may lead to subtle deviations from the assumption of universal fitness effects, strain-specific immune responses require to invoke fitness effects that are tied to a specific host at a specific time. For this reason, we relied on data collected at the very early stages of infection when specific immune responses have not yet been mounted. It is important to note, however, that host differences will lead to less convergent evolution making the observed patterns of sharing even more inconsistent with a neutral model of evolution.

## Methods

### Data and identification of shared mutations

For all of the individuals from Keele et al. (2008) and Li et al. (2010), a sequence sample and a consensus sequence are available. Only the sequences from the individuals in which the infection was the result of a single founder virus were used. These were all aligned to a reference sequence HIV-1 NL-43 (Di Giallonardo et al., 2013). Mutations in two populations are considered shared if they map to the same position in the reference genome and have the same base in the consensus sequence that changed into the same mutated base. Deletions and insertions were not considered.

Sequences from this dataset where mutations carrying an APOBEC signature are removed using hypermut (Rose and Korber, 2000) were acquired from Elena Giorgi. Any mutations that do not occur in these sequences, but are present in the original sequences are considered to be APOBEC-mediated mutations.

### Simulations of sequence evolution

Simulations of sequence evolution were adapted from Lee et al. (2010). The first step of the sequence evolution simulation is the initialization, which consists of ancestor sequence generation and the initialization of the fitness table. The sequence generation is the same as in Lee et al. and results in a random sequence, *s*, with the same length N and base distribution as the *env*-gene. For this sequence, a fitness table can then be created. This is a table of size N by 4, where each entry is the fitness effect of each possible base at every position. The entries for the ancestor sequence are set to 1, the rest is filled up with random draws from the mutational fitness effects distribution.

During the simulation, each sequence in every generation generates offspring according to Poisson(*R*_0 *_ *ƒ*), where *ƒ* is the fitness, as

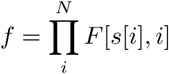

This is the product of the entries in the fitness table for all positions in the sequence. Every new offspring sequence acquires new mutations according to mutation rate *μ* and substitution matrix M. Optionally, the G-to-A mutation rate can be increased x-fold to simulate APOBEC-mediated G-to-A hypermutation. The population is capped to 10000 sequences, as this is the effective population size of HIV. Once the number of sequences exceeds this cap, 10000 sequences are randomly sampled from the population and used as the next generation.

In order to recreate the dataset, a separate simulation is performed for each individual with the same initial sequence and fitness table. After each generation, a sample is taken from the simulation with the same size as the number of sequences available for the individual. The number of unique mutations in this sample is then counted. If this matches the data, the simulations are stopped and this sample is used for analysis. If this number exceeds the amount present in the data, the previous sample is also compared and the sample with the closest amount of mutations is used.

Since all individual simulations are, apart from the initial generation of ancestor sequence and fitness table, completely independent, these simulations can be run in parallel, reducing the required time per simulation drastically. The simulations were implemented in python, parallelized with MPI4PY (Dalcín et al., 2005) and run on a cluster.

## ABC-SMC

The ABC-SMC framework (standing for Approximate Bayesian Computation using Sequential Markov Chains, Toni et al. (2009)) was used for simultaneous model selection and parameter estimation of the MFED. For this, we implemented an SMC procedure in python.

The fitting procedure starts with equal probabilities for all models. Simulations are then performed for all models with random parameters according to their prior distribution. The priors were uniform distributions from zero to one for all parameters except λ (0, 2) in the exponential beneficials models, *μ* (– 1,1) in the log-normal models and *b*_*b*_ (1,2) in the 5 spikes model. In the first iteration, a set of parameters (a ‘particle’) is sampled and a simulation is run with these parameters. If the distance between the summary statistics of this simulation and the data is smaller than ϵ_1_, the particle is retained. Once 1000 particles have been accepted, a weight is assigned to all accepted particles (see Toni et al. (2009)).

In the next generations, a particle is sampled from the previous iteration using the assigned weights. The particles are then perturbed according to Gaussian kernel and a simulation is run with these parameters. Again, the distance between the summary statistics of this simulation and the data is calculated. If this is smaller than *ϵ*_*i*_ (with *i* the iteration), the particle is retained and once 1000 particles have been accepted, the weights are calculated.

We used tolerances *ϵ*_*i*_ = [2.2,1.3, 0.8,0.7, 0.6,0.5, 0.4], and a Gaussian kernel with *σ* equal to the average distance between accepted particles in the model divided by two. We used a normalized Euclidean distance to calculate the distance between simulations and the data, where the summary statistics are divided by the corresponding summary statistics of the data, after which the distance to a vector of ones is calculated.

The model probabilities are directly calculated from the set of accepted parameters (the number of accepted particles for model *x*/1000). It is possible for a model to ‘die out’ during the SMC if none of the particles for this model are accepted. This indicates an exceedingly low model probability.

For the parameter estimation we did a Gaussian kernel density estimation for the accepted parameters in the SMC in the last iteration for each remaining model. From this, we determined the highest density point - which was used as the parameter estimate - and the 95% highest density intervals.

### Summary Statistics

In total 14 summary statistics were defined to calculate the distance between simulations in data. The majority of them are based on shared mutations, which are defined as a mutation that occurs in the same position with the same from-and to-base in two independent populations.

Five of the summary statistics are based on the average degree of of sharing of the mutations in an individual, which is defined as the sum of all mutations, weighted by the amount of other populations said mutations appear in (the weight for a unique mutation is thus 0, the weight of a mutation occurring in 3 populations total is 2), divided by the total amount of mutations in the viral population. This number can be calculated for all inidividuals in a dataset. The resulting distribution of values can be characterized by the mean, the median, the difference between mean and median, the variance and the area under the cumulative curve of the sorted degrees of sharing per viral population.

Where the degree of sharing characterizes sharing from the point of view of the inidividual, the occurrence of mutations is a similar measure, this time from the point of view of the mutations. It measures in how many populations each mutation is present. These numbers are then sorted and categorized. For the summary statistics, the following five categories are used: the number of mutations occurring only once (singletons), the amount occurring in exactly two populations, those occurring three to five times, five to ten times and finally the number of mutations occurring in more than 10 populations.

The remaining four summary statistics are individual-based, and three serve as a proxy to match the number of generations to the data. They are calculated for each population separately, and averaged for the summary statistic. A first measure is the fraction unmutated sequences, the percentage of sequences in the sample that are identical to the founder. For the simulations the founder is known, for the data the consensus sequence is assumed to be the same as the founder sequence. The fraction unmutated is then used to also calculate a selection index, the percentage of sites in the sample that are mutated in at least one sequence, divided by the percentage of mutated sequences in the sample (which is 1-%unmutated). Lastly, the average inter-sequence hamming distance is calculated per viral population, which, according to Lee et al. (2010), is directly related to the time since infection in a neutral model.

In order to match the G-to-A mutation rate increase as well, the fraction of G-to-A mutations in the sample was calculated per inidividual. The mean of this distribution was then used as the last summary statistic.

